# Reproductive capacity evolves in response to ecology through common developmental mechanisms in Hawai’ian *Drosophila*

**DOI:** 10.1101/470898

**Authors:** Didem P. Sarikaya, Samuel H. Church, Laura P. Lagomarsino, Karl N. Magnacca, Steven Montgomery, Donald K. Price, Kenneth Y. Kaneshiro, Cassandra G. Extavour

## Abstract

Lifetime reproductive capacity, or the total number of offspring that an individual can give rise to in its lifetime, is a fitness component critical to the evolutionary process. In insects, female reproductive capacity is largely determined by the number of ovarioles, the egg-producing subunits of the ovary. Recent work has provided insights into the genetic and environmental control of ovariole number in *Drosophila melanogaster*. However, whether regulatory mechanisms discovered under laboratory conditions also explain evolutionary variation in natural populations is an outstanding question. Here we report, for the first time, insights into the mechanisms regulating ovariole number and its evolution among Hawai’ian *Drosophila*, a large adaptive radiation of fruit flies in which the highest and lowest ovariole numbers of the genus have evolved within 25 million years. Using phylogenetic comparative methods, we show that ovariole number variation among Hawai’ian *Drosophila* is best explained by adaptation to specific oviposition substrates. Further, we show that evolution of oviposition on ephemeral egg-laying substrates is linked to changes the allometric relationship between body size and ovariole number. Finally, we provide evidence that the developmental mechanism principally responsible for controlling ovariole number in *D. melanogaster* also regulates ovariole number in natural populations of Hawai’ian drosophilids. By integrating ecology, organismal growth, and cell behavior during development to understand the evolution of ovariole number, this work connects the ultimate and proximate mechanisms of evolutionary change in reproductive capacity.

## Introduction

Reproductive capacity is an important life history trait that directly influences fitness by determining how many offspring an individual can leave behind. There is a wide range in potential fecundity across species (1, 2), which is often interpreted as trade-offs with presumed ecological and developmental constraints. Trade-offs have been invoked to explain patterns of egg-laying in animals, where total fecundity can correlate negatively with egg mass, clutch size or lifespan (3-10), and positively with body size (11-13). In addition to these hypothesized physical or growth-related constraints, life history parameters including predation risk, environmental variability, host specialization and levels of parental care have been proposed to influence evolutionary change in fecundity (1, 14-17), suggesting that this trait could represent a complex intersection between ecology and physiology. However, few studies have addressed how female reproductive capacity evolves in response to ecology, and how these pressures manifest as different phenotypes through changes in development.

In insects, female reproductive capacity is strongly influenced by the number of egg-producing structures called ovarioles (1, 18-23). Ovariole number is species-specific and genetically determined (24, 25). Most insects have limited intraspecific variation in ovariole number, and the effect of ovariole number on fecundity has been observed by comparing mean ovariole numbers within or between species. In many insects, including beetles, fruit flies, and aphids, ovariole number is positively correlated with fecundity between and within species (1, 21-23). For example, *Drosophila melanogaster* strains with naturally occurring or genetically manipulated higher ovariole numbers both show increased fecundity (18, 26). While physiological traits like egg production rate may also play an important role in determining reproductive capacity (27), these can be difficult to assess in laboratory settings where egg-laying conditions may not be suitable for some insects. In contrast, ovariole number has served as a proxy for reproductive capacity for decades (18), as it is a quantitative trait that can be easily measured from field and laboratory samples.

Ovariole number is established during larval and pupal stages (20), and can be affected by environmental conditions during this phase of development, including nutrition and temperature (24, 28, 29). During larval development, a specific group of cells called terminal filament cells (TFCs) form stacks called terminal filaments (TFs) that serve as the beginning point of each ovariole (30-33). Developmental mechanisms of ovariole number evolution are best characterized in species of the African *melanogaster* subgroup of *Drosophila*, where average ovariole number ranges from 43 to 18 per female (1, 34), and ovariole number differences result primarily from changes in TFC number (29, 35). Ovariole number is highly polygenic and regulated by pleiotropic genes (25), thus offering an opportunity to study the evolution of a complex quantitative trait in response to different environments.

Major shifts in ovariole number have been attributed to aspects of life history. Ovoviviparity, where females oviposit first instar larvae, is often correlated with reduced ovariole number (16), suggesting that increased parental investment is linked to reduced fecundity as observed in other animals (17). The stability of the environment and the predictability of egg-laying substrates may influence evolution of ovariole number, as more stable environments or abundant substrates are correlated with higher ovariole number, and species occupying unpredictable environments or scarce substrates tend to have lower ovariole numbers (15, 36). In the well-studied *Drosophila melanogaster* subgroup, previous studies have suggested that reproductive strategies and ovariole number evolve in response to oviposition or larval nutrition substrate (35-37). Most *melanogaster* subgroup species are generalists that oviposit on a variety of decaying fruits, and mean ovariole number in this subgroup ranges from 43 to 18 per female (1, 34). In contrast, *D. erecta* and *D. sechellia* are specialists on *Pandanus* fruit and the toxic *Morinda* fruit, respectively (38, 39), and *D. sechellia* has the lowest reported ovariole number of the group (1). This reduction in ovariole number has been hypothesized to be the result of increased egg size as an adaptation to feeding on the toxic *Morinda* (40), or to be due to lower insulin signaling levels evolved in response to the relatively constant nutritional input provided by substrate specialization (35). Reviewing data on oviposition behavior in *melanogaster* subgroup species, Lachaise (37) proposed that the high ovariole number observed in the generalists *D. melanogaster* and *D. simulans* may be driven by the frequent oviposition opportunities available to these species, as they oviposit on most decaying fruit. However, the *melanogaster* subgroup is not well-suited for a broader understanding of ovariole number evolution, as most species share similar oviposition substrates (i.e. rotting fruit) and there are few independent instances of evolution of specialists.

In contrast, Hawai’ian *Drosophila* have evolved to specialize on a variety of oviposition substrates, including decaying flowers, leaves, fungi, sap fluxes, and bark of native plants, and eggs of native spiders (41). Moreover, these flies exhibit the most extreme interspecies range of ovariole number reported in the genus, ranging from two to 101 per ovary (42). Hawai’ian *Drosophila* have undergone rapid island radiation from a common ancestor in the last 25 million years, leading to over 1000 extant species (43-45). The high species diversity of Hawai’ian *Drosophila* is spread across five monophyletic species groups that share genetic, morphological and ecological similarities, and rely on different oviposition substrates (44, 46-48), as follows (Figure 1): *Scaptomyza* are small species that primarily lay eggs on leaves or flowers. Picture wing (PW) species are larger species with striking pigment patterns on their wings (49). PW species primarily lay eggs on decaying bark or branches of native trees, though some specialize on sap fluxes (41). Modified mouthpart (MM) species, which have male-specific modifications on mouthparts used during mating (50), have the largest range of egg-laying substrates, specializing on decaying leaves, fungi, sap or bark (51). Haleakala species are darkly pigmented flies that only lay eggs on native fungi. Lastly, most antopocerus-modified tarsus-ciliated tarsus (AMC) species are leaf breeders, though there are a few exceptions that have evolved bark-breeding (44).

**Figure 1.**
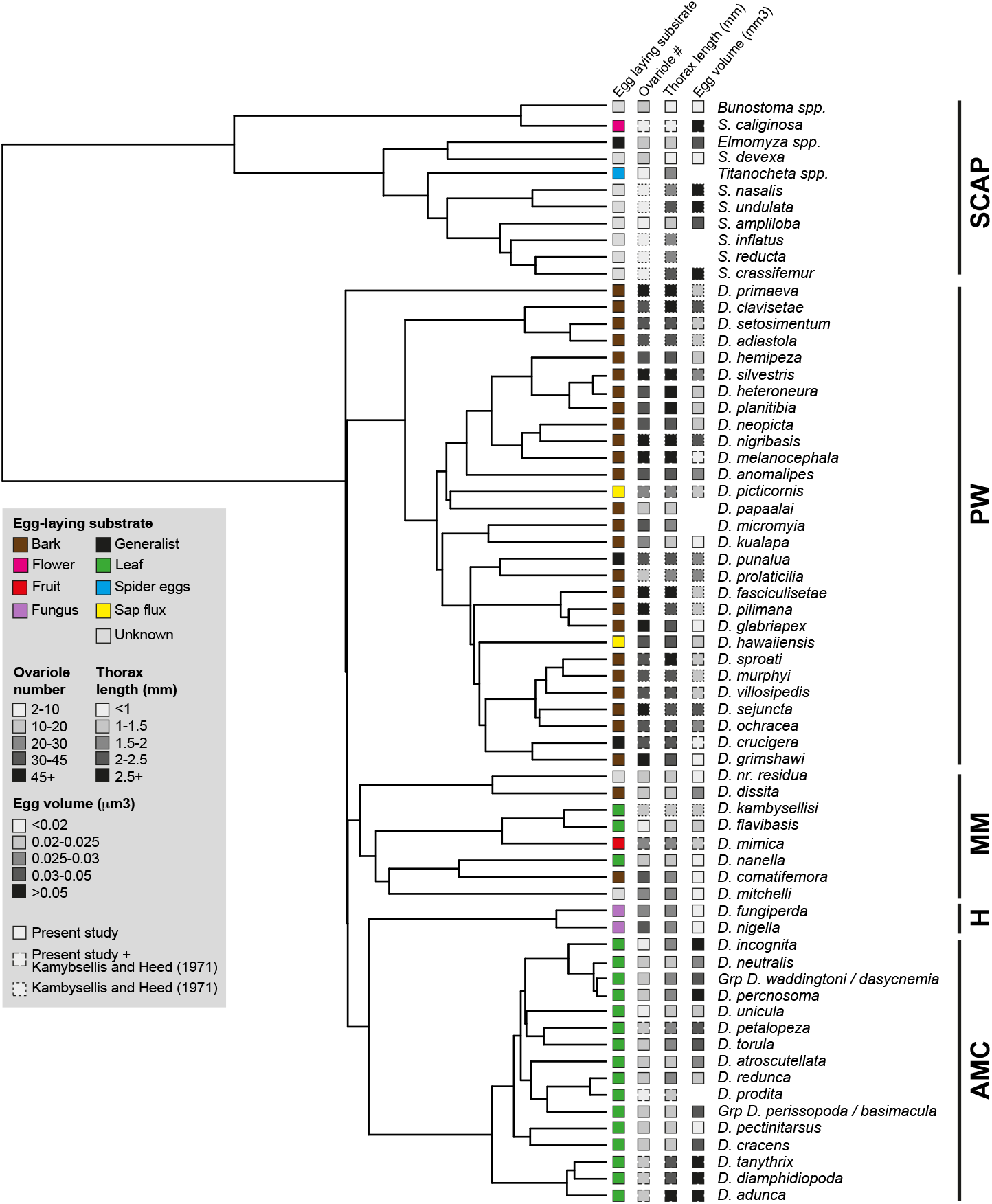
Reproductive and ecological traits of Hawai’ian *Drosophila* in phylogenetic context. Compiled adult life history traits (greyscale gradients) collected herein and by Kambysellis and Heed (42) are mapped on a phylogeny of Hawai’ian *Drosophila* constructed from available mitochondrial and nuclear genes. Egg-laying substrate of each species is indicated by colored boxes: bark (brown), generalist (black), sap flux (yellow), leaf (green), fungus (purple), fruit (red), spider eggs (blue), flowers (pink), and unknown (gray). Boxes with solid outlines denote data collected in the present study; boxes with four notches denote data represented in our data and those of Kambysellis and Heed (42); boxes with dotted outline denote data represented only in Kambysellis and Heed (42). Missing boxes indicate data points that were either not previously reported (42) or that we were unable to obtain from field-caught samples. Black lines at right delineate the five major groups of Hawai’ian *Drosophila* as follows: SCAP =*Scaptomyza;* PW = picture wing; MM = modified mouthparts; H = Haleakala; AMC = antopocerus-modified tarsus-ciliated tarsus.

Ovariole number is highest in the PW species (up to 202 per female), and lowest in Scaptomyza and AMC species (as few as 2 per female) (42). Dramatic differences in ovariole number between species have been hypothesized to be a result of shifts between their varied oviposition substrates (42, 51). Other studies have posited that the divergent ovariole number observed in Hawai’ian *Drosophila* may be a result of r-K evolution (42), given the surface area of decaying trees, and the predictability of this substrate in the field (36), is greater than that of other oviposition substrates (51, 52). However, the studies supporting these hypotheses primarily sampled PW species, and used phylogenies that have since been substantially improved upon in more recent studies that include expanded taxon sampling and additional loci (44, 46, 48, 53).

To investigate the linked effects of ecology and development underlying ovariole number evolution in Hawai’ian *Drosophila*, we conducted phylogenetic comparative analyses of life history traits from 60 species, and comparative development analyses from ten species using both wild-caught flies and laboratory strains. Our results identify potential mechanisms of evolutionary change in ovariole number operating at three levels of biological organization. First, we found that evolutionary shifts in ecological niche could predict the dramatic differences in ovariole number in Hawai’ian *Drosophila*. Second, whether adult body size was coupled with ovariole number or egg volume differed between species groups with different oviposition substrates, suggesting that the allometric growth relationships between these traits evolves dynamically. Finally, we found that changes in ovariole number from two to 60 per individual can be explained by changes in total TFC number, suggesting that ovariole number diversity evolves through the same developmental mechanism, regardless of the specific ecological constraints or selective pressures.

## Results and Discussion

### Adult reproductive traits of Hawai‗ian Drosophila

We measured three major adult traits relevant to reproductive capacity (body size, ovariole number and egg volume), from field-collected females, lab-reared F1 offspring of field-collected females, and females from laboratory strains (Figure 1; Table S1). Species identities of field-collected females were assigned based on morphological keys or DNA barcoding (Tables S2, S3). All traits ranged over an order of magnitude within Hawai’ian *Drosophila:* body size ranged from 0.71mm for *S. devexa* to 3.12mm for *D. melanocephala*, ovariole number per female ranged from 2 for *S. caliginosa* to 88.5 for *D. melanocephala*, and egg volume ranged from 0.01 um^3^ for *Bunostoma spp*. group (*S. palmae/S. anomala*) to 0.2um^3^ for *D. adunca*, highlighting the diversity of life history traits in Hawai’ian *Drosophila*.

Within the *melanogaster* subgroup species, species-specific differences in ovariole number are largely heritable (25, 54, 55). To test whether this is also the case in Hawai’ian *Drosophila*, we compared ovariole number of wild-caught females and their lab-reared F1 offspring, across five species with different egg-laying substrates. We observed no significant differences between the ovariole numbers of these two generations regardless of natural substrate (Figure S1), indicating that species-specific differences in ovariole number are also strongly genetically determined in Hawai’ian *Drosophila*.

### Larval ecology influences ovariole number evolution

A previous study based almost exclusively on picture wing species proposed that evolutionary shifts in larval ecology had driven ovariole number diversification in these flies (51). To test this hypothesis across the major groups of Hawai’ian *Drosophila*, we compared the fit of evolutionary models of ovariole number that accounted for ecologically driven evolution, to those that did not. Our dataset included both specialist species that oviposit on one of bark, sap flux, leaf, fungus, fruit, flower or spider-eggs, as well as generalist species that oviposit on multiple decaying substrates (Figure S2). We compared the fit of five models to our data, two of which ((i) Brownian Motion, BM, and (ii) an Ornstein Uhlenbeck model with a shared optimum for all species, OU1) do not take into account the oviposition substrate, and three of which were nested ecological models based on alternative methods of substrate classification: (iii) OU2 assumed two states, bark breeders and all other species, to test previous suggestions that bark-breeding may drive evolution of ovariole number (51, 52); (iv) OU3 assumed three states, *Scaptomyza* specialists on spider eggs and flowers, bark-breeders, and species using any other substrate, to test proposals that substrates influence ovariole number evolution because of their differences in carrying capacity and field predictability (36, 42); and (v) OU8 categorized each oviposition substrate separately. These five models were fit over 100 trees sampled from the posterior distribution of a Bayesian phylogenetic analysis to account for phylogenetic uncertainty.

We found that models accounting for larval ecology explained the ovariole number diversification in Hawai’ian *Drosophila* (Table 1) better than those that did not. Comparing the three ecological models, we found that the three-state model (OU3), which accounted for both bark breeders and *Scaptomyza* specialists, was supported as the best-fit model across a majority of trees for ovariole number (ΔAICc > 2 as compared to OU2 and OU8 models; Table S4). Estimated theta values for the OU3 model showed that bark breeders have more ovarioles than species that oviposit on other substrates, suggesting that evolution of higher ovariole numbers accompanied the transition to bark breeding from likely non-bark breeding ancestors (Fig. 2A,B, Table S5), consistent with earlier hypotheses (51, 52). In contrast, *Scaptomyza* species may have experienced a dramatic decrease in ovariole number as they independently specialized on spider eggs and flowers (Fig. 2B). Taken together, our results suggest that shifts in oviposition substrate may have contributed to the evolution of diverse ovariole numbers in this group, not only for picture wing flies as predicted previously (51), but across the adaptive radiation of Hawai’ian *Drosophila*.

**Table 1.** Comparison of AICc and weighted AICc values for models testing the relationship between oviposition substrate and ovariole number. Values are for model fit of Brownian motion (BM) and Ornstein-Uhlenbeck with one optimum (OU1) or with multiple optima (OUM) with different combinations of oviposition substrate categories, calculated with the R package **OUwie** v.1.48 (75). Oviposition substrates were categorized as follows: OU2 categorizes species that lay eggs on bark and non-bark; OU3 categorizes species into bark-breeder, spider egg/flower breeder, and other; and OU8 categorizes each species according to the eight oviposition substrates represented (bark, flower, spider egg, fruit, leaf, generalist, fungus, sap flux). Models were tested over 1000 posterior distribution BEAST trees using nuclear and mitochondrial gene sequences. Bold indicates the best supported model.

| | AICc | ΔAICc | w(AIC) |
| --- | --- | --- | --- |
| BM | 86.26 | 5.91 | 0.04 |
| OU1 | 88.41 | 8.06 | 0.01 |
| OU2 | 84.84 | 4.49 | 0.1 |
| **OU3** | **80.35** | **0** | **0.77** |
| OU8 | 84.42 | 4.07 | 0.08 |

**Figure 2.**
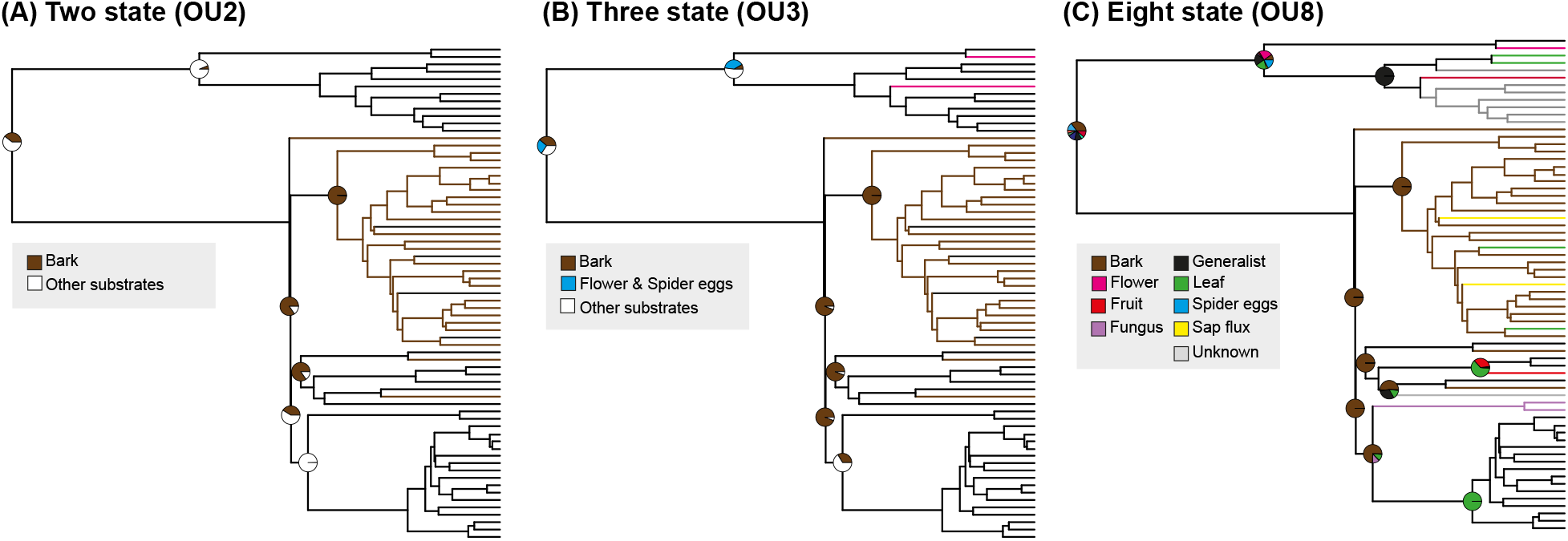
Different ecological states tested for OU analysis. (A) A two-state model (OU2) of bark-breeders (brown) and non-bark breeders (white). (B) Three-state model (OU3) that codes bark-breeders (brown), spider egg and flower breeders (blue), and other oviposition substrates (white). (C) Eight-state model (OU8) that codes each egg-laying substrate separately, color coded as in Figure 1. Pie charts show the maximum likelihood ancestral state estimates at each node, calculated with the rayDISC function in the R package **corHMM**, v. 1.18 (74).

In African drosophilids and tephritid *Dacus* flies, generalist species that oviposit on a variety of egg-laying substrates have higher fecundity than specialists (1, 22, 37). Moreover, specialist species of African and Central American *Drosophila* species are more fit in the presence of host-specific compounds (40, 56-58), some of which are toxic to other species of *Drosophila*. For example, *D. sechellia* is best reared on lab media supplemented with *Morinda* fruit (40), while *D. pachea* cannot be reared in laboratory conditions without supplementing media with sterols from its host cactus (59). Egg-laying substrates for Hawai’ian *Drosophila* have divergent chemical cues and fungal populations (60). Hawai’ian *Drosophila* often lay few eggs on unsupplemented laboratory food (see Supplemental Information), but do not change ovariole number when reared on this food (Figure S1). We therefore speculate that specific substrate components may not only allow females to distinguish between hosts for oviposition, but also may contribute to species- and substrate-specific egg laying behavior in Hawai’ian *Drosophila*.

### Evolution of specialist habitats changes allometry of reproductive traits

The range of Hawai’ian *Drosophila* body sizes is greater than that of other members of the genus, spanning an order of magnitude (Table S1). To determine whether changes in allometric growth might underlie reproductive trait evolution, we analyzed the allometric ratio of such traits using a phylogenetic least squares (PGLS) analysis and thorax volume (thorax length^3^) as a proxy for body size. We found that across all Hawai’ian *Drosophila*, thorax volume was significantly positively correlated with both ovariole number (Figure 3A; Table 2; Table S6) and egg volume (Figure 3B; Table 2; Table S6).However, individual species groups show differences in trends for allometric ratios of reproductive traits. In PW and MM species, body size is correlated positively with ovariole number (Figure 3A1, A2), but not with egg volume (Figure 3B1, B2). In contrast, AMC and *Scaptomyza* species have a positive correlation with body size and egg volume (Figure 3B3, B4), but not ovariole number (Figure 3A3, A4). For PW, MM, and AMC, there is a negative correlation between ovariole number and proportional egg size (Table S2; Figure S3B-D), and there is a negative correlation between ovariole number and egg volume in AMC and *Scaptomyza* (Table 2; Figure S3I-J).

**Table 2.**
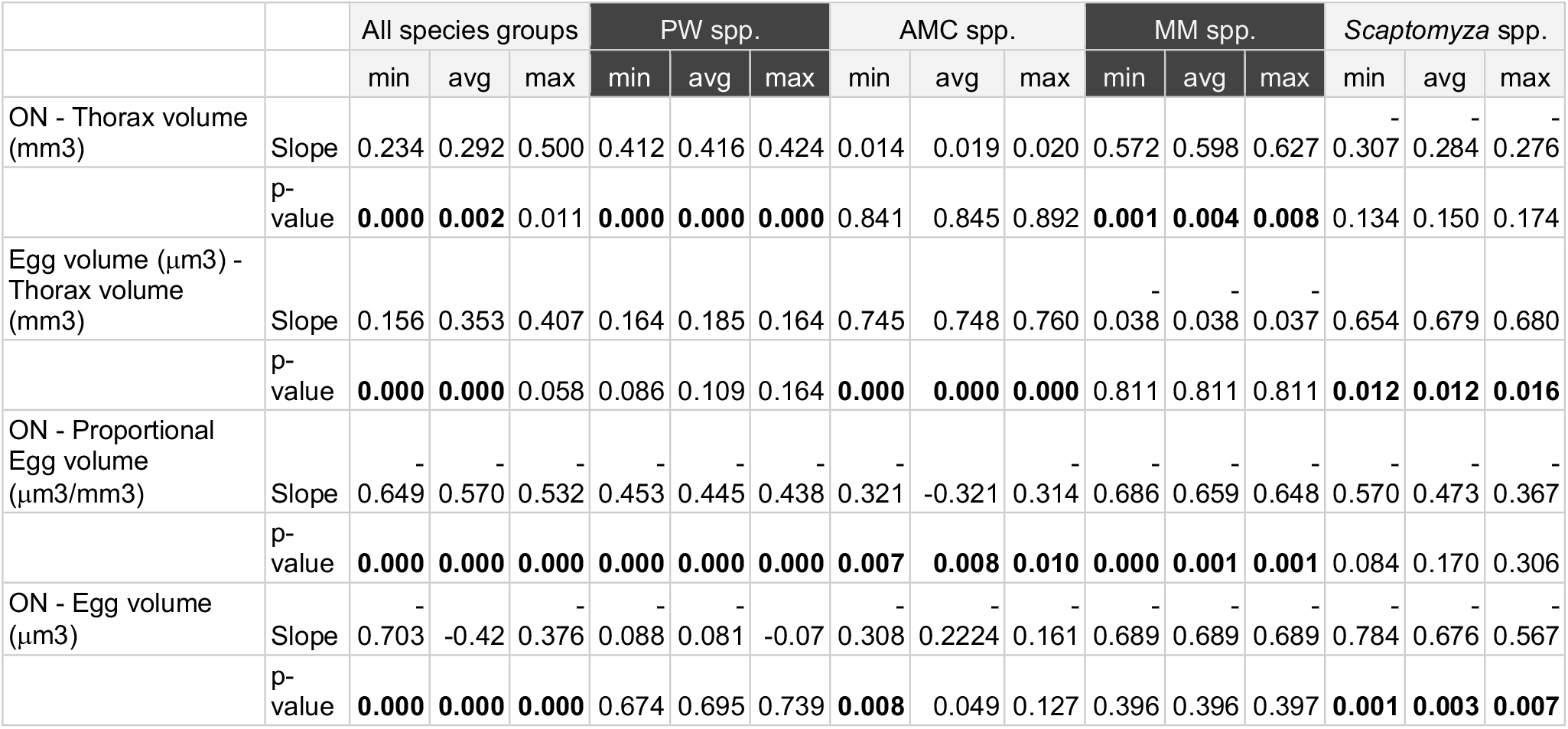
Phylogenetic Generalized Least Squares (PGLS) analysis of adult reproductive traits in Hawai’ian *Drosophila*. PGLS analysis of relationships between ovariole number and thorax volume (mm^3^), egg volume (μm^3^) and thorax volume, and ovariole number and proportional egg volume (μm^3^/mm^3^) are listed. Regression analyses were performed with the R package **nlme** v.3.1-121 (76) on 100 trees from a BEAST posterior distribution using nuclear and mitochondrial genes, and the minimum, average, and maximum slope and p-value for the analysis is included in the table. P-values below 0.01 are indicated in bold.

**Figure 3.**
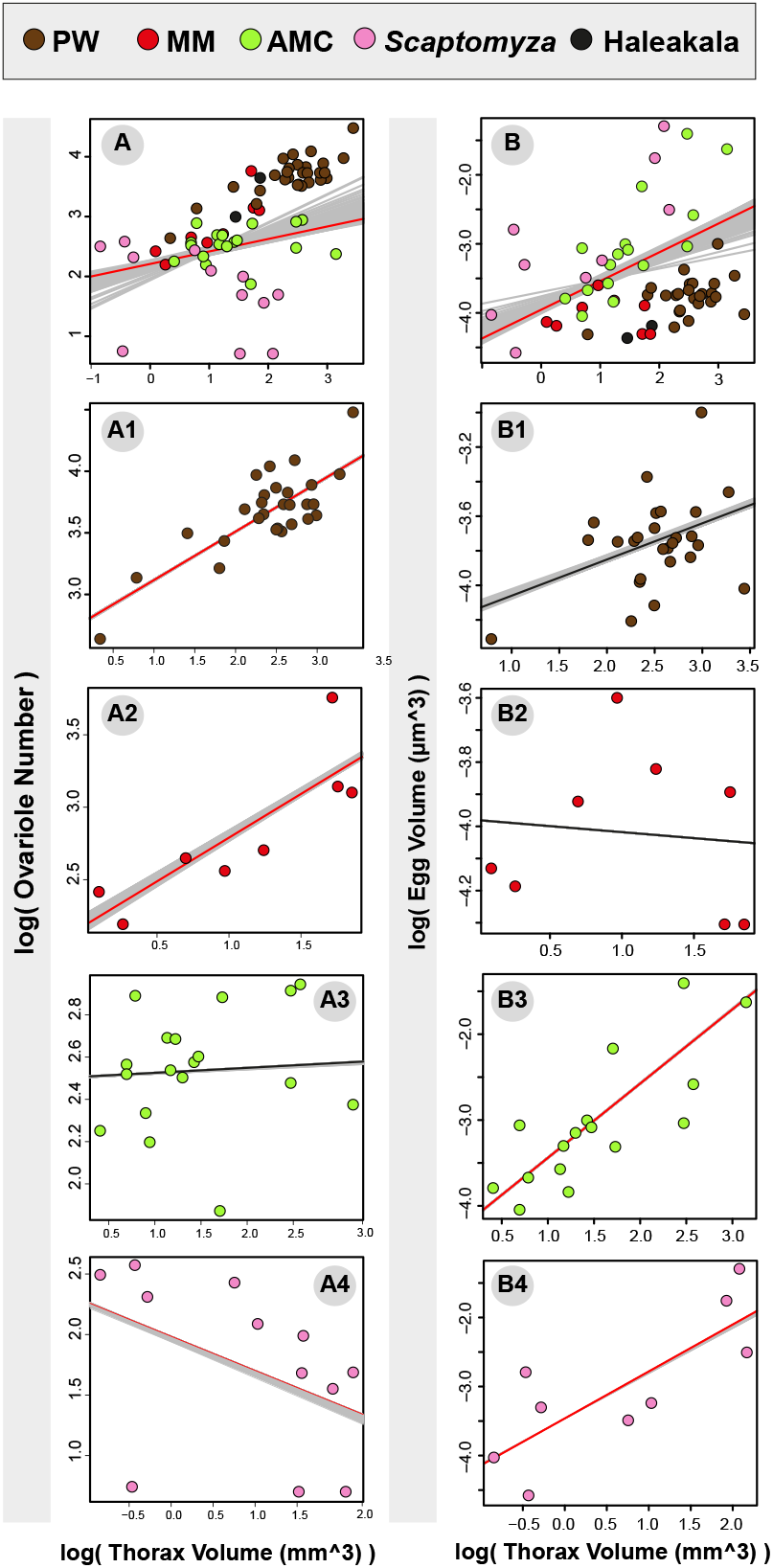
Allometric relationship between life history traits in Hawai’ian *Drosophila*. Scatter plots of log transformed adult measurements with phylogenetically transformed trend lines generated by averaging runs from PGLS analysis across 100 posterior distribution BEAST trees, performed with the R package **nlme** v.3.1-121 (76). Trend line of the consensus tree is denoted in red when there was a significant relationship between the two traits, and black when PGLS analysis did not support a significant relationship (Table 2). (A, A1-A4) Ovariole number plotted against thorax volume (mm^3^) in (A) all specimens, (A1) PW, (A2) MM, (A3) AMC, and (A4) *Scaptomyza*. (B, B1-B4) Egg volume (μm^3^) plotted against thorax volume (mm^3^) in (B) all specimens, (B1) PW, (B2) MM, (B3) AMC, and (B4) *Scaptomyza*.

We note that these trends are associated with differences in life history strategies between groups. PW and MM group species, in which ovariole number increases with increasing body size (Figure 3A1, A2), lay eggs in abundant and varied substrates (41): PW are primarily bark breeders that oviposit eggs in clutches of up to 100 eggs (42), and MM group species can occupy a wide range of oviposition preferences, including bark, leaf, fruit, fungus, and sap flux (41). In contrast, species of AMC and *Scaptomyza*, in which ovariole number and body size are decoupled (Figure 3A3, A4), have independently evolved use of substrates with low carrying capacity: AMC group species are primarily leaf breeders, reproducing on damp leaves in the forest bed, while the oviposition substrates of *Scaptomyza* species include ephemeral substrates, such as flowers, spider eggs and fresh leaves, many of which are not occupied by other Hawai’ian *Drosophila* species groups (41). In sum, while a positive correlation between body size and fecundity is commonly posited in egg-laying animals (11, 13), we did not find universal support for this trend across Hawai’ian *Drosophila*. This is, however, consistent with previous studies on Diptera, wherein trends toward higher fecundity or ovariole number in larger animals observed within species (11) contrast with between-species differences in ovariole number that do not always correlate with body size (22, 37, 61).

### Larval ovary somatic cell number determines ovariole number

We previously identified two developmental mechanisms that can alter ovariole number during development: changes in TFC number per TF and change in total TFC number (29). To determine whether the same developmental mechanisms that regulate ovariole number in laboratory populations, also underlie the evolution of ovariole number in natural populations, we measured TF and TFC numbers in the developing larval ovaries of Hawai’ian *Drosophila*. Our analysis of 12 species representing four of the major Hawai’ian *Drosophila* species groups showed that even over a range of ovariole numbers spanning an order of magnitude (Figure 4; Table S7), larval TF number essentially corresponded to adult ovariole number (Table S8). Although TFC number per TF varied somewhat between species (Figure 4A; Table S7), PGLS analysis showed no correlation between TFC number per TF and total TF number (Table 3). In contrast, average total TFC number was strongly positively correlated with TF number (Table 3; Figure 4B; Table S7), suggesting that, as in laboratory populations of *D. melanogaster*, changes in TFC number underlie ovariole number evolution in Hawai’ian *Drosophila*.

**Figure 4.**
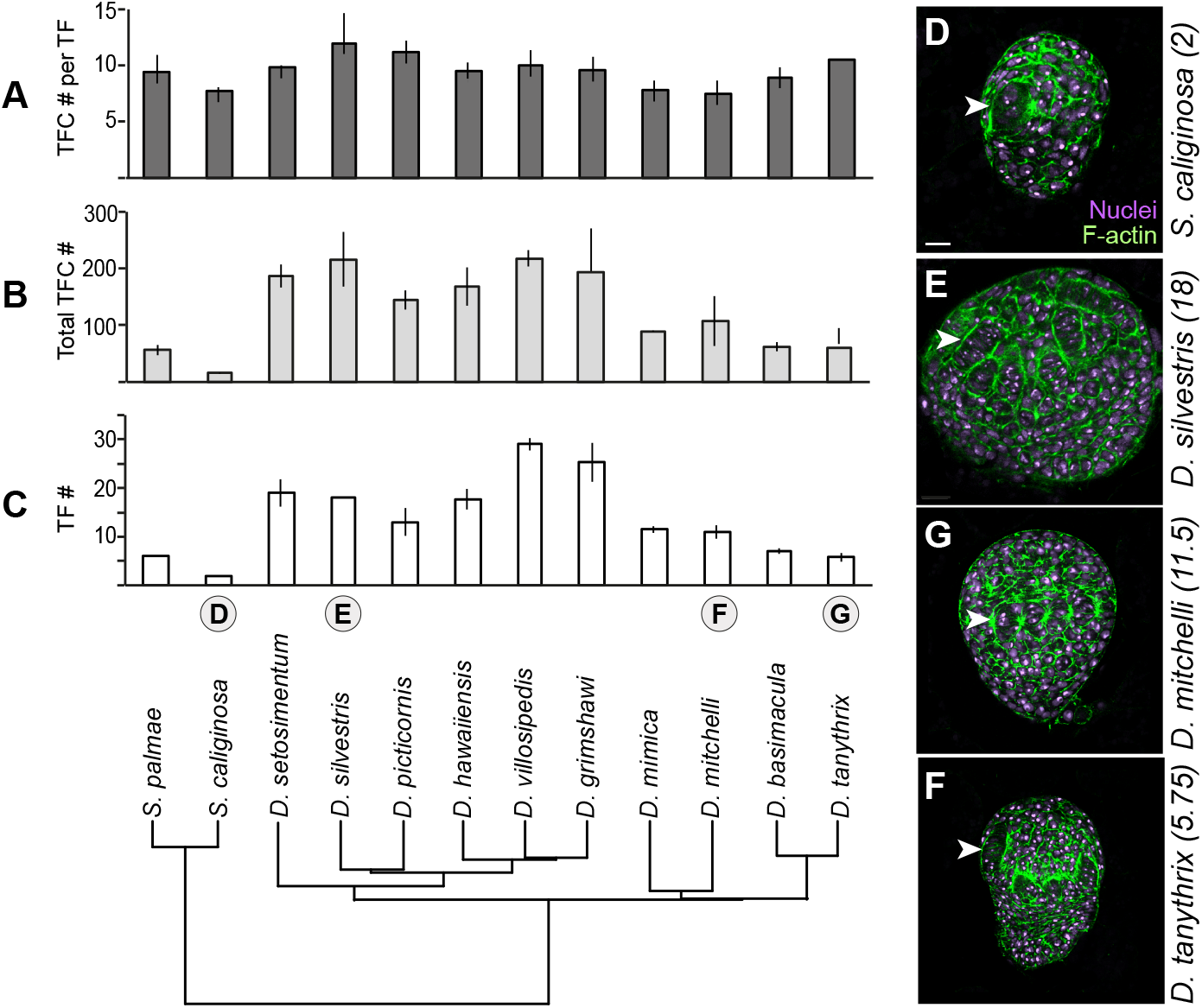
Terminal filament cell (TFC) number predicts terminal filament (TF) number in Hawai’ian Drosophilids. (A-C) Bar graphs for (A) TFC number per TF, (B) total TFC number, and (C) TF number per larval ovary representing the mean and standard deviation, as well as the phylogenetic relationship between the species shown (bottom). (D-F) Late third instar larval ovaries stained for nuclei (purple) and F-actin (green) for (D) *S. caliginosa* (flower breeder), (E) D. *silvestris* (bark breeder), (G) *D. mitchelli* (egg-laying substrate unknown), and (F) *D. tanythrix* (leaf breeder). Numbers in parentheses beside species names indicate mean ovariole number per ovary (Tables S7, S8). White arrowheads indicate TF structures in the ovary.

**Table 3.** Phylogenetic Generalized Least Squares (PGLS) analysis of larval ovarian measurements in Hawai’ian *Drosophila*. Relationships between TF number and TFC number per TF, TF number and total TFC number, and total TFC number and TFC number per TF are listed. Regression analyses were performed with the R package **nlme** v.3.1-121 (76) on 100 trees from a BEAST posterior distribution using nuclear and mitochondrial genes, and the minimum, average, and maximum slope and p-value for the analysis is included in the table. P-values below 0.01 are indicated in bold.

|  |  | min | avg | max |
| --- | --- | --- | --- | --- |
| TF # - TFC # per TF | Slope | 0.320 | 0.744 | 1.728 |
|  | p-value | 0.199 | 0.376 | 0.647 |
| TF # - Total TFC # | Slope | 0.873 | 0.873 | 0.873 |
|  | p-value | **0.000** | **0.000** | **0.000** |
| TFC # per TF - Total TFC # | Slope | 0.097 | 0.097 | 0.097 |
|  | p-value | 0.059 | 0.059 | 0.059 |

The developmental mechanism underlying ovariole number evolution is particularly interesting in light of the allometric changes in Hawai’ian *Drosophila* species groups. There has been some debate as to whether allometry constrains or facilitates adaptive evolution (62-64). In Hawai’ian *Drosophila*, the allometric relationship between two important female reproductive traits, ovariole number and egg size, was coupled to body size in different groups in different ways: when ovariole number was coupled with body size, egg size was not, and vice versa (Figure 3). These trends were associated with abundant versus scarce egg-laying substrates respectively (Figure 1). While the phenotypic integration of ovariole number and egg volume appears tightly regulated across insects (65), the coupling of ovariole number to body size appears more flexible in Hawai’ian *Drosophila*, suggesting that in this context, heritable changes in allometry may contribute to adaptive evolution.

Ovariole number is regulated by both by intrinsic and extrinsic growth factors, including Hippo signaling, ecdysone and insulin-like peptides, all of which can also regulate body size (26, 35, 66-68). Thus, we propose that the mechanistic basis for evolutionary change of ovariole number on different substrates, may be changes in the relative influence of nutritionally regulated circulating growth factors on the one hand, and cell-autonomous growth on the other hand, on ovarian development during larval and pupal stages. For example, we speculate that on certain substrates, the larval ovary may become less sensitive to nutritionally-mediated growth factors by evolving lower expression levels of growth factor receptors, and relying more on tissue-specific growth factors, which could include local insulin release or cell proliferation pathways such as Hippo signaling.

Taken together, we found that highly divergent ovariole number, and by proxy female reproductive capacity, have evolved together with changes in egg-laying substrate across Hawai’ian *Drosophila*. Moreover, this remarkable adaptive radiation is linked to evolutionary changes in a key reproductive trait that is regulated by variation in the same developmental mechanisms operating in the model species *D. melanogaster*.

## Materials and Methods

Hawai’ian *Drosophila* were collected (69) at the Koke’e State Park and Kui’a NAR on Kauai, West Maui Watershed Reserve, Makawao Forest Reserve, and Waikamoi Nature Preserve on Maui, and the Volcanoes National Park and Upper Waiakea Forest Reserve on Hawai’i island. Field-caught flies were brought back to the lab for species identification and phenotyping of adult and larval characters. Measurements of adult ovariole number, larval TF and TFC number were performed as previously described (29). Mature egg size and adult body size were quantified from white light micrographs of eggs and adult thoraces using ImageJ. See Supplementary Information for detailed methods.

We combined sequence data for 18 genes reported in four previous studies (44, 46, 48, 53) from GenBank with additional newly identified mitochondrial sequences (Table S9), and used the concatenated sequences to generate trees in RAxML v8.2.3 (70). Phylogenetic relationships and divergence time estimates were inferred in a Bayesian framework in BEAST v. 2.3.2 (71, 72). All phylogenetic comparative analyses and corresponding figures were computed in R version 3.2.0 (73).

We used reported ecological information about Hawai’ian *Drosophila* to code oviposition site (41), calculated ancestral states for each of these character codings with BEAST using the *rayDISC* function in the R package **corHMM**, v.1.18 (74), mapped the most likely ecological state at each node, and pruned the resulting tree to include only tips with ovariole number data. The fit of different models of trait evolution was assessed on the pruned trees in **OUwie** v. 1.48 (75). See Supplementary Information for detailed methods and custom scripts.

## Acknowledgements

This work was supported by National Institutes of Health grant number 1R01 HD073499 to CGE; National Science Foundation (NSF) Doctoral Dissertation Improvement Grant number DEB-1209570, a post-graduate scholarship from the Natural Sciences and Engineering Research Council of Canada (NSERC) and a pre-doctoral fellowship from the Fonds de recherché du Québec— Santé (FRQS) to DPS; NSF Graduate Training Fellowship to SHC; and NSF Postdoctoral Research Fellowship in Biology Grant number 1523880 to LPL.

